# Functional interplay between H2B ubiquitylation and H2A.Z deposition

**DOI:** 10.64898/2026.04.22.720199

**Authors:** Younus A. Bhat, Ivano Mocavini, Erin O’Donnell, Song Tan, Oliver J. Rando, Craig L. Peterson

**Author notes:** These authors contributed equally to this work (Younus A. Bhat, Ivano Mocavini).

## Abstract

The Bre1 ubiquitin conjugating enzyme catalyzes the monoubiquitylation of histone H2B-K120/K123 at promoter proximal nucleosomes, contributing to transcriptional regulation. These same nucleosomes are targeted by the yeast SWR1C chromatin remodeling enzyme that deposits the histone variant H2A.Z. Yeast strains that lack both Bre1 and Swr1 are inviable, indicating that together they contribute to an essential cell function. Interestingly, H2B-K123ub and H2A.Z levels are anticorrelated, and recent biochemical studies suggest a model in which H2B-K123ub inhibits SWR1C activity by blocking access to the nucleosome acidic patch. Here, we exploit recombinant nucleosomes to show that the H2A.Z deposition activity of SWR1C is strongly inhibited by H2B-K123ub. Surprisingly, the loss of H2B-K123ub does not lead to significant changes in the genomic organization of H2A.Z in cells grown in normal media, but we find that H2B-K123ub is required for a re-localization of H2A.Z from promoters to coding regions during replication stress. Together, these data indicate a complex functional interplay between H2B ubiquitylation and H2A.Z deposition.

## Introduction

Eukaryotic genomes are assembled into a nucleoprotein structure called chromatin which at the most basic level consists of nucleosomes in which ∼147bp of DNA wrapped onto an octamer of the core histone proteins, H2A, H2B, H3, and H4^1^. Nucleosome assembly hinders the accessibility of DNA to the machineries that control gene transcription, chromosome replication, and genome stability pathways. Consequently, several mechanisms exist that regulate chromatin dynamics, facilitating nuclear processes. These mechanisms include histone posttranslational modifications (PTMs), ATP-dependent chromatin remodelers, and the incorporation of histone variants^2,3^. Furthermore, these factors that regulate chromatin dynamics often exhibit functional interplay; for instance, histone lysine acetylation can facilitate the recruitment of chromatin remodelers to genomic loci, as many remodelers harbor bromodomain-containing subunits that directly recognize this PTM ^4^.

The monoubiquitylation of histone H2B at lysine 120 in humans or lysine 123 in yeast is a PTM associated with actively transcribed genes, and it is required for the methylation of histone H3 at lysines 4 and 79 at promotor proximal nucleosomes and coding regions, respectively^5,6^. H2B-K120/K123ub is catalyzed by the Bre1 ubiquitin ligase, and this PTM is subject to de-ubiquitylation by a set of ubiquitin-specific proteases, such as yeast USP8 and USP10. The dynamic cycles of ubiquitylation and de-ubiquitylation of H2B leads to an enrichment of H2B-K120/123ub at the promoter proximal, +1 nucleosome (for an excellent discussion see ^6^). Although loss of H2B-K123ub does not have a large impact on the steady state RNA pool in yeast^7^, studies have demonstrated disruption of RNA polymerase II dynamics during gene transcription due to loss of either this PTM or the resulting loss of histone H3 methylations^6^. H2B-K123ub also limits both cryptic transcription and antisense transcripts, likely due to its cooperation with the FACT complex to re-assemble nucleosomes following passage of RNAPII ^8,9,10^. In addition to its role in transcription, H2B-K123ub plays an important role during DNA replication in response to replication stress in yeast^11, 12^. Yeast strains that lack H2B-K123ub (e.g. *bre1* mutants) are sensitive to the replication stress agent, hydroxyurea (HU), and such mutants show defects in replication fork stalling in response to HU treatment^11,12^. These changes in fork progression appear to be due to a defect in nucleosome assembly behind the advancing replication fork, and it has been suggested that H2B-K123ub may also regulate nucleosome assembly during transcriptional elongation^7,12^.

The large, multi-subunit SWR1C enzyme is a yeast member of the INO80 subfamily of ATP-dependent chromatin remodeling enzymes, and it catalyzes the stepwise replacement of nucleosomal histone H2A/H2B dimers with histone variant H2A.Z/H2B dimers^13–16^. SWR1C is targeted to nucleosomes that flank the promoter-proximal, nucleosome-depleted region (NDR), and incorporation of H2A.Z at the +1 nucleosome facilitates the kinetics of gene activation^17,18^. In addition, H2A.Z plays a global role in yeast transcription when cells lack the nuclear exosome^19^. Yeast strains that lack both SWR1C and the Bre1 ubiquitin ligase show synthetic lethality, indicating that they function in parallel, redundant pathways for an essential function^20^. Notably, H2A.Z levels anti-correlate with H2B-K120ub; for instance, at the +1 nucleosome, H2B-K123ub is enriched at the NDR proximal face while H2A.Z is located at the NDR distal surface^21^. H2A.Z incorporation has been shown to inhibit the ubiquitin ligase activity of Bre1 and the human homology, RNF20/40^22^, but whether H2B-K123ub regulates SWR1C activity is not known.

The nucleosome surface contains a solvent-exposed acidic ‘patch’ that serves as a binding site for many nonhistone proteins, including chromatin remodelers and histone modifying enzymes^23^. Recently, we found that the SWR1C remodeler requires the nucleosome acidic patch, and evidence suggested that the conserved Swc5 subunit makes a direct contact with this domain^24^. Interestingly, H2B-K120/123ub is located in close proximity to the acidic patch, and recent structural data indicate that this mark can impede access to the acidic patch by nonhistone proteins^25^. Here we test if H2B-K123ub impacts the H2A.Z deposition activity of SWR1C, using defined, recombinant nucleosome substrates. We find that H2B-K123ub has a strong negative impact on SWR1C activity, but this mark has little impact on the nucleosome sliding activity of the RSC remodeler. Surprisingly, loss of H2B-K123ub in vivo has little impact on the genome-wide distribution of H2A.Z under normal growth conditions, suggesting that the inhibition observed in vitro may not contribute significantly to the anticorrelation between H2A.Z and H2B-K123ub enrichment at promoter nucleosomes. In contrast, we find that H2B-K123ub is required for the accumulation of H2A.Z at stalled replication forks in yeast cells exposed to replication stress.

## Results

### H2B-K123ub inhibits SWR1C dimer exchange activity

To investigate the impact of H2B ubiquitylation on dimer exchange by SWR1C, center-positioned nucleosomes were assembled with recombinant histone octamers containing *Xenopus laevis* H3/H4 tetramers and either wildtype *Saccharomyces cerevisiae* H2A/H2B (AB) dimers or *Saccharomyces cerevisiae* H2A/H2B dimers containing a monoubiquitin conjugated to an engineered cysteine residue on H2B-K123, which we will refer to as H2B-K123ub (Figure 1B). The DNA template for nucleosome reconstitutions was a 207-bp fragment containing a ‘‘601’’ nucleosome positioning sequence and symmetric 31 bp linker DNAs (31N31). Following reconstitution, H2B-K123ub nucleosomes were observed to migrate noticeably slower compared to unmodified nucleosomes on native PAGE gels (Figure 1C). These nucleosomes were employed in gel-based assays that monitor deposition of H2A.Z (Figure 1A). In one assay, nucleosome substrates were incubated with excess SWR1C, ATP, and free H2A.Z/H2B dimers in which H2B contains a 3xFLAG tag at its C-terminus. Reaction products are separated on native PAGE, and formation of the heterotypic (AB/ZB) and homotypic (ZB/ZB) nucleosomal products are detected by their reduced gel migration due to the 3xFLAG tag on H2B, as visualized by SYBR staining. Alternatively, the free H2A.Z/H2B dimers were labelled with Cy5, allowing H2A.Z deposition to be monitored by the gain of Cy5 fluorescence within the nucleosomal product.

**Figure 1.**
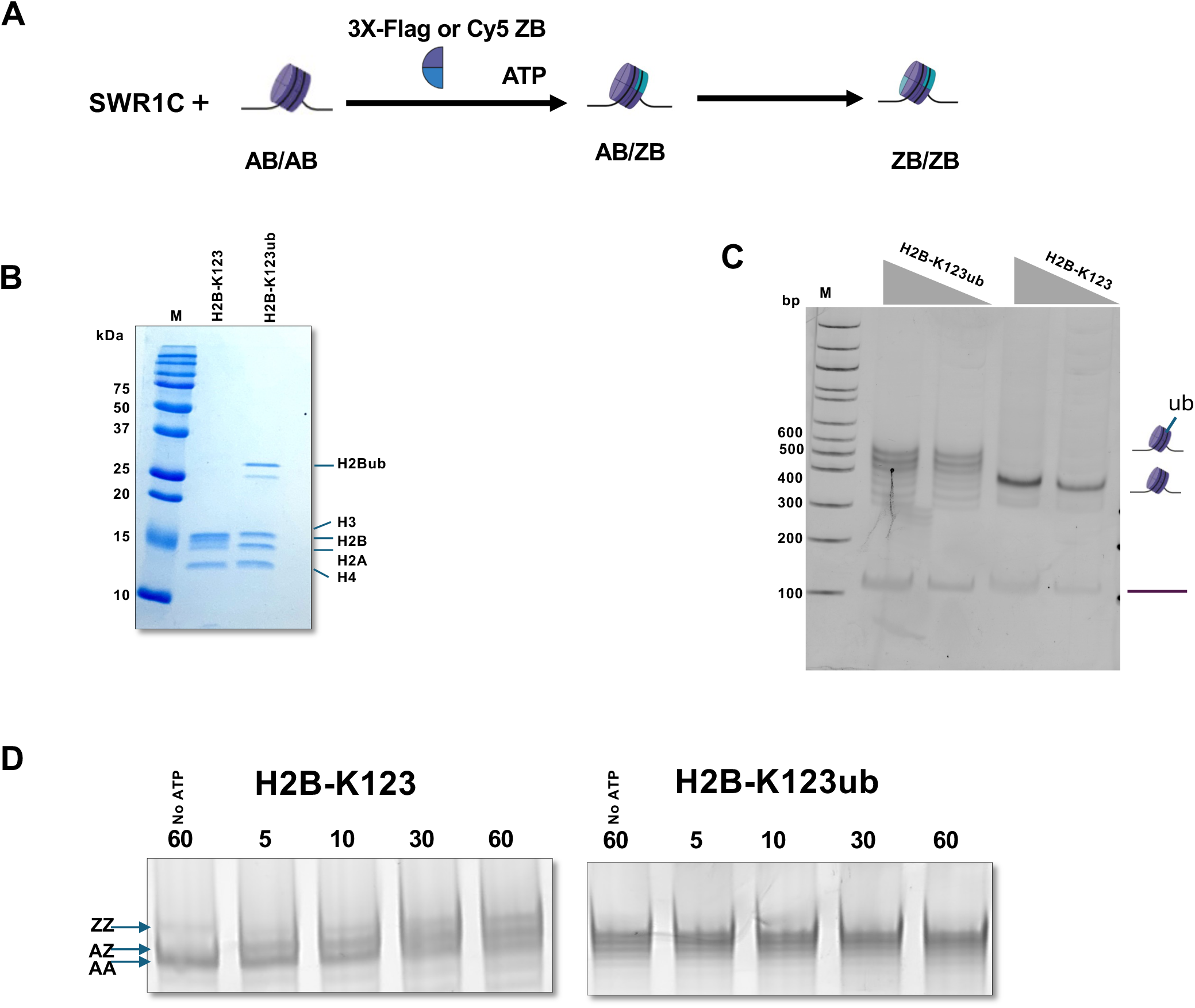
H2B-K123ub inhibits SWR1C-mediated dimer exchange activity. a) cartoon representation of SWR1C mediated dimer exchange activity; b) H2B-K123 and H2B-K123ub octamers electrophoresed on a 15% SDS-PAGE, confirming presence of ubiquitin on H2B-K123; c) H2B-K123 and H2B-K123ub nucleosomes electrophoresed on 4% native PAGE; d) SWR1C-mediated dimer exchange activity on H2B-K123 and H2B-K123ub nucleosomes with 3X-Flag tagged ZB dimers as the co-substrate.

Addition of SWR1C to unmodified nucleosomes led to the time- and ATP-dependent accumulation of slower migrating nucleosomal species when H2A.Z/H2B-3xFLAG dimers were used in the exchange reaction (Figure 1D). In contrast, the banding pattern of the H2B-K123ub nucleosomes remained unchanged in the SWR1C reactions (Figure 1D). Notably, dimer exchange would involve loss of the H2B-K123ub-marked dimers, which is expected to lead to faster migrating species, even taking into account the changes in migration due to the 3xFLAG-tagged dimers. When Cy5-labelled H2A.Z/H2B dimers were used in the exchange reactions, SWR1C addition promoted an accumulation of Cy5-labelled nucleosomes, consistent with productive dimer exchange on the unmodified nucleosomes (Figure S1). In contrast, reactions containing H2B-K123ub did not lead to the expected faster migrating species, and there was little ATP-dependent increase in Cy5 fluorescence (Figure S1). Together, these data indicate that H2B-K123ub strongly inhibits the dimer exchange activity of SWR1C.

These data support the view that H2B-K123ub inhibits SWR1C activity. Alternatively, the data might suggest that the integrity of H2B-K123 is key, rather than the ubiquitin mark. To test this possibility, we performed dimer exchange assays with nucleosomes reconstituted with octamers containing either H2B-K123, H2B-K123C, H2B-K123A, H2B-K123R, or H2B-K123W. As shown in Figure 2, none of the H2B substitution derivatives had a significant impact on SWR1C dimer exchange activity, using H2A.Z/H2B-3xFlag dimers in the assay. Thus, the inhibition of SWR1C appears to require the ubiquitin modification, rather than the integrity of H2B-K123.

**Figure 2.**
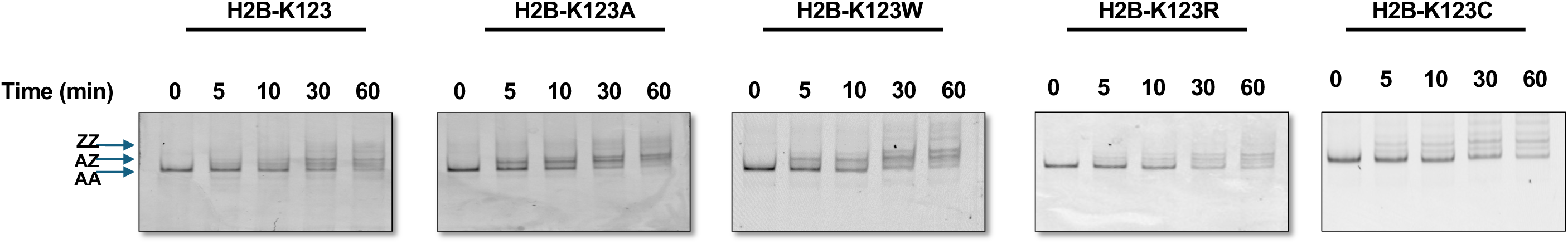
SWR1C mediated dimer exchange activity is not impacted by amino acid substitutions at H2B-K123. Histone octamers with different H2B-K123 substitution derivatives were reconstituted into Cy3-labelled, end-positioned nucleosomes.

The RSC remodeler is a member of the SWI/SNF subfamily of remodelers, and it catalyzes the ATP-dependent mobilization of nucleosomes in cis, as well as histone octamer eviction^2^. Like other members of the SWI/SNF family, RSC subunits make contact with the nucleosomal acidic patch, and these interactions contribute to nucleosome binding^26,27^. We employed a restriction enzyme accessibility assay to monitor the ability of RSC to mobilize 31N31 nucleosomes that contain H2B-K123ub (Figure 3A). In the absence of RSC, a *Hha*I site located near the center of the positioned nucleosome is occluded, with very little cleavage even after 60 minutes of incubation (Figure 3B). Addition of RSC and ATP led to significant HhaI cleavage even after 5’ of incubation with the unmodified nucleosome (Figure 3B,C). In contrast, H2B-K123ub had no detectable impact on RSC activity (Figure 3B,C). Thus, H2B-K123ub has a selective inhibitory impact on members of two different families of remodelers.

**Figure 3.**
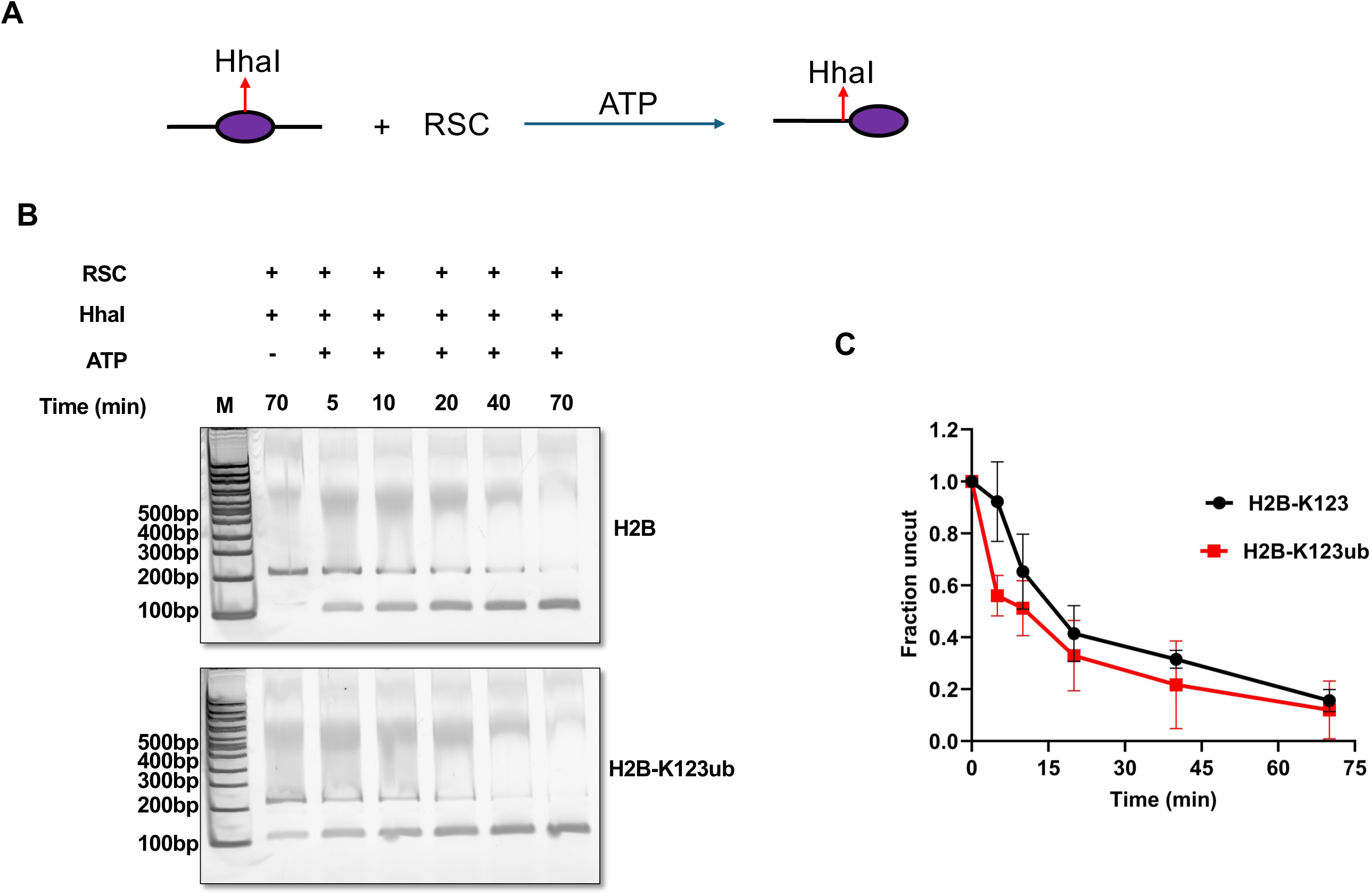
RSC mediated restriction accessibility assay on H2B-K123 and H2B-K123ub nucleosomes. a) Cartoon representation of restriction accessibility assay using RSC; b) Native-PAGE showing restriction accessibility assay using H2B-K123 (upper gel) and H2B-K123ub (lower gel) nucleosomes; c) plot showing fraction uncut with time in presence of RSC, ATP, and *Hha*I. Band intensity for uncut and cut was measured using image J software and ratio was plotted against time.

### Interplay between H2A.Z deposition and H2B-K123ub in yeast

To investigate the impact of H2B ubiquitylation on H2A.Z deposition in yeast, isogenic wildtype and *bre1*Δ** strains were created that harbored a FLAG-tagged allele of *HTZ1* which encodes yeast H2A.Z (Figure S2A). Bre1 is the ubiquitin ligase for H2B-K123, and consequently the *bre1*Δ** strain lacks H2B-K123ub, as well as H3-K4me3 which relies on H2B ubiquitylation (Figure S2A). Loss of Bre1 had no impact on either the total cellular levels of H2A.Z or the fraction bound to chromatin (Figure S2B). To investigate the impact of the *bre1*Δ** on the genome-wide distribution pattern of H2A.Z, chromatin immunoprecipitation analyses were performed. In our wild-type control, we recapitulated the previously reported localization of H2A.Z, with maximal enrichment at genic +1 nucleosomes and more modest enrichment at −1 nucleosomes (Figure 4A,B). Furthermore, H2A.Z levels were generally low over gene coding regions, as expected (Figure 4B,C). Surprisingly, the distribution of H2A.Z was not changed significantly in the *bre1* mutant, with levels at both promoter-proximal nucleosomes and gene coding regions remaining nearly identical to those observed in the wildtype strain (Figure 4B,C).

**Figure 4.**
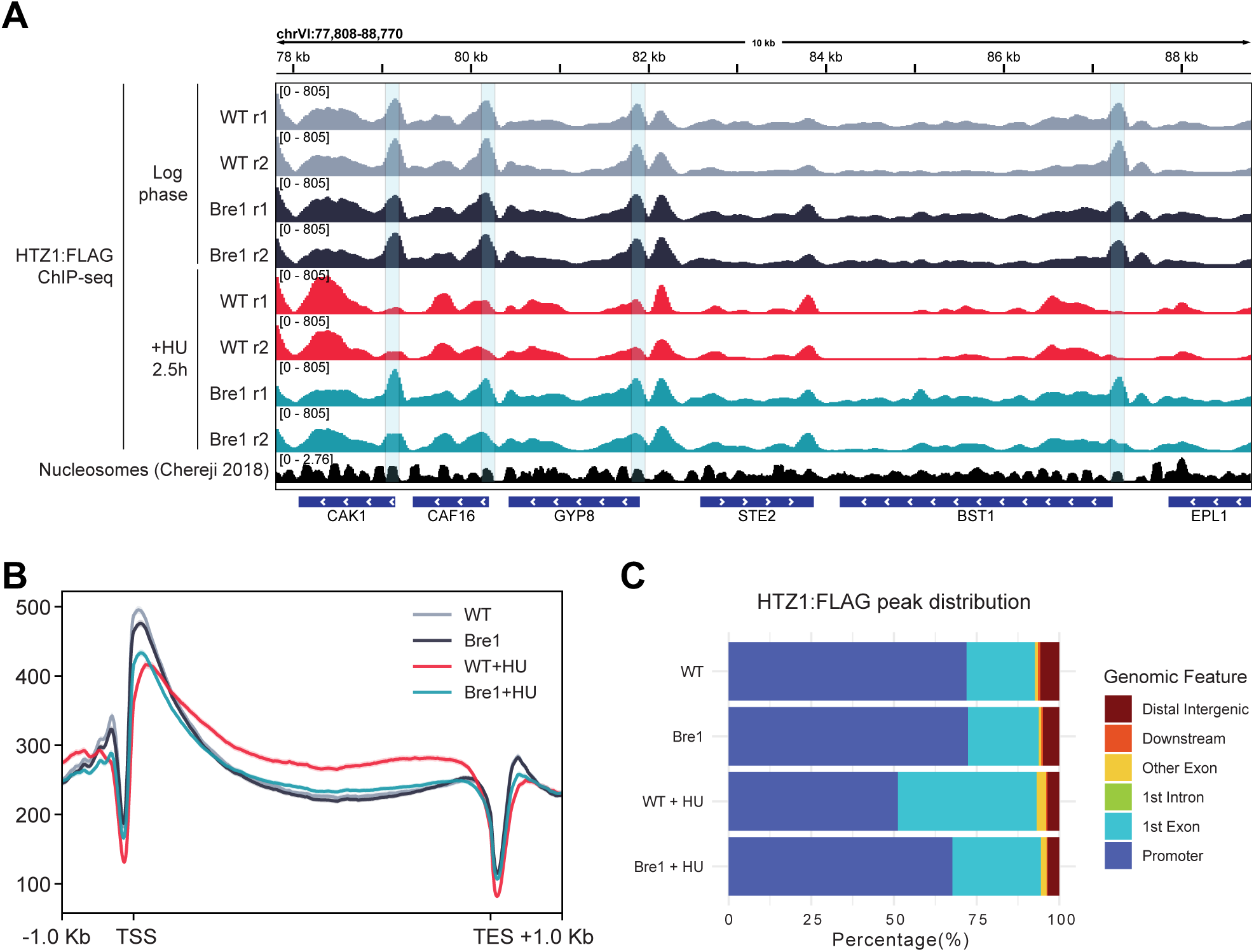
Genome-wide Htz1 distribution in the absence of H2B-K123ub. A) Genome view of *HTZ1:FLAG* ChIP-seq tracks of individual replicates of WT and *bre1*Δ**, in normal growing conditions and upon HU treatment. Nucleosomal occupancy data from Chereji et al., 2018 is also added as a reference^42^. Shaded boxes highlight the +1 nucleosomes, over which Htz1 signal is lost in WT cells upon HU treatment; b) metagene plot of Htz1 average levels over yeast genes; c) Genomic annotation of Htz1 peaks.

Given that H2B-K123ub plays an important role during DNA replication in response to replication stress, the genome-wide distribution of H2A.Z was also monitored following a 2h 30m treatment with hydroxyurea where cells accumulate in early to mid S phase. In HU-treated wildtype cells, H2A.Z was depleted from promoter-proximal nucleosomes, and increased levels of H2A.Z were observed over gene coding regions (Figure 4A–C). Importantly, these changes were primarily observed at genes replicated early in S phase compared to those replicating late in S phase, indicating that the changes in H2A.Z distribution were associated with replication progression (Figure 5). In contrast, H2A.Z distribution remained relatively unchanged in the HU-treated *bre1* mutant, even though the *bre1* strain showed a similar S phase accumulation compared to the wildtype (Figures 4,5 and Figure S3). Thus, these data indicate that H2A-K123ub is required for a re-distribution of H2A.Z at newly replicated genes in cells undergoing replication stress.

**Figure 5.**
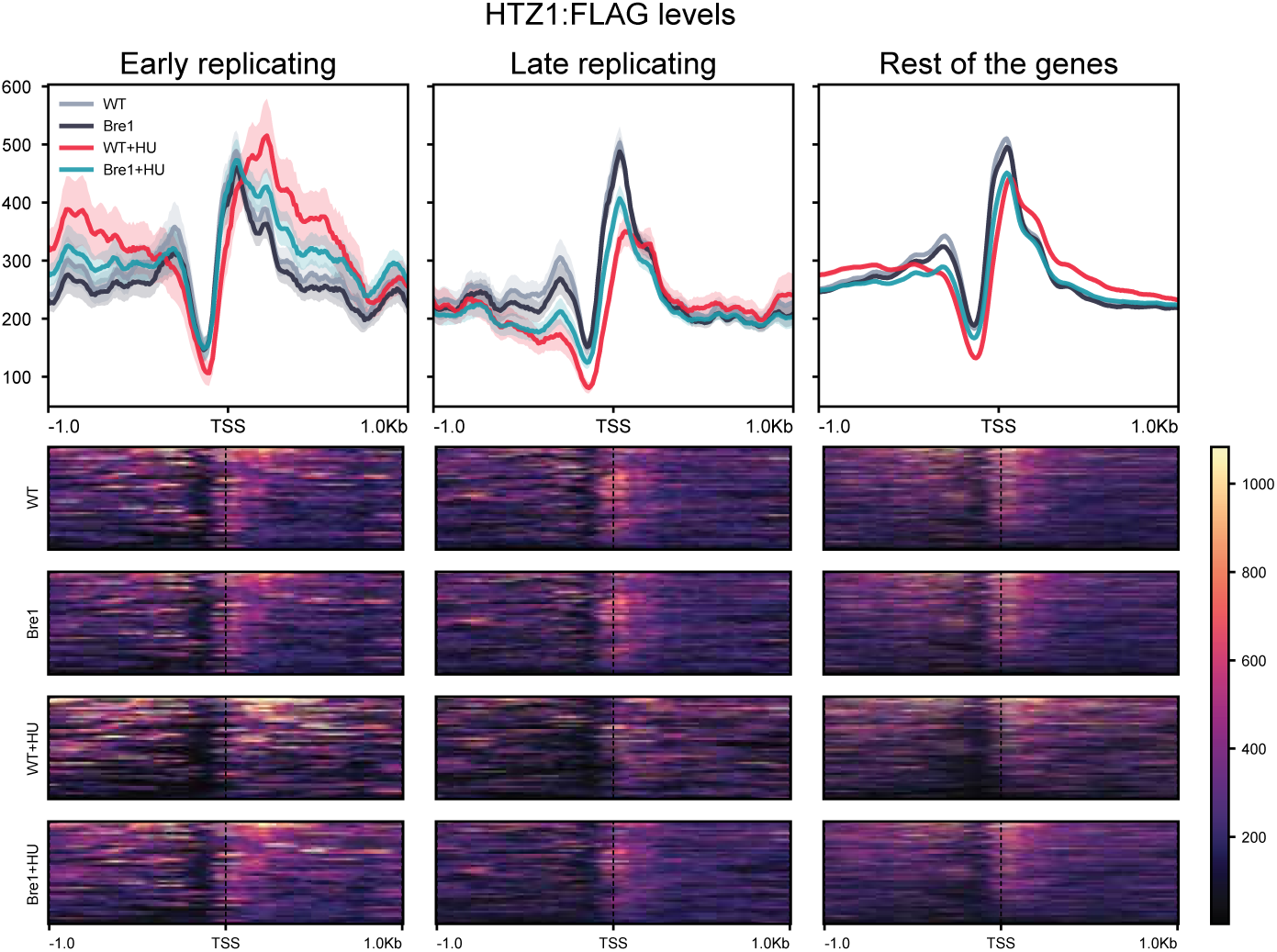
Replication-dependent Htz1 redistribution is impacted in *bre1*Δ** cells. a) metagene profile (top) and single-locus heatmap (bottom) Htz1:FLAG ChIP-seq signal over early replicating, late replicating, and rest of the genes. Each line represents the average signal of two biological replicates, the shaded area represents the standard error.

## Discussion

An emerging feature of ATP-dependent chromatin remodeling enzymes is that they each make functional interactions with the solvent-exposed acidic patch on their nucleosomal substrates^26–33^. Recently, we reported that SWR1C requires both nucleosomal acidic patches to perform even a single cycle of H2A.Z deposition, and the Swc5 subunit was identified as a putative acidic patch interacting subunit.^24^ Structural studies have suggested that the binding of nonhistone proteins to the acidic patch might be generally inhibited by the ubiquitylation of H2B-K123, as this mark lies in close proximity on the nucleosomal surface^25^. Indeed, our recent cryoEM model of a Swc5-nucleosome complex suggested that its interactions with the acidic patch might be blocked by ubiquitylation of H2B-K123^24^. Here we tested this idea directly with recombinant nucleosomal substrates, finding that H2B-K123ub inhibits the H2A.Z deposition activity of SWR1C, while having no detectable impact on the ability of the RSC remodeler to mobilize nucleosomes in cis.

Although remodelers generally require interactions with the nucleosomal acidic patch, regulation by H2B ubiquitinylation is remodeler specific. For instance, H2B-K123 ubiquitylation does not inhibit the Chd1 remodeler, but in fact this modification increases the remodeling rate of Chd1 by 2- to 3-fold^34^. The Chd1 remodeler utilizes the energy of ATP hydrolysis to mobilize nucleosomes in cis, and like SWR1C, the optimal rate of this nucleosome mobilization reaction requires an intact nucleosomal acidic patch. However, in the case of Chd1, H2B-K123 ubiquitylation does not block interactions with the acidic patch, but rather the ubiquitin group appears to stabilize an initial, unwrapped DNA state, leading two the increased rate of nucleosome mobilization^34^.

Members of the SWI/SNF subfamily of remodelers, such as yeast RSC or mammalian BAF complexes, interact with both nucleosomal acidic patches, ‘clamping’ the nucleosome via two distinct binding domains^26, 35,36^. Thus, as expected, alteration of the acidic patch has a dramatic impact on the remodeling activity of these enzymes^37^. Although the dependency on an intact acidic patch is shared between SWI/SNF family members and SWR1C, we found that the remodeling activity of RSC was not impacted by H2B-K123ub. Likewise, work with mammalian BAF complexes showed that the cBAF and PBAF complexes were not inhibited by H2B ubiquitinylation^37^. Surprisingly, the ncBAF complex was inhibited by H2B ubiquitinylation, even though this BAF assembly is believed to only contact one of the two acidic patches^37^. Thus, the relationship between acidic patch interactions and H2B ubiquitinylation is not straightforward.

H2B-K123ub (K120 in mammals) is a conserved mark that is enriched at the promoter proximal, +1 nucleosome, a site also enriched for the histone variant H2A.Z ^6,7^. High resolution, ChIP-exo studies found that H2B-K123ub and H2A.Z are anti-correlated at the +1 nucleosome, with each mark enriched at opposite faces of the nucleosome^21^. Such anticorrelation is consistent with prior biochemical studies demonstrating that the yeast Bre1 ligase and the mammalian RNF2/40 ubiquitinylation machinery are less active on an H2A.Z nucleosome ^22^, as well as data shown here where H2B-K123ub hinders H2A.Z deposition. It was anticipated that loss of H2B-K123ub might lead to higher levels of H2A.Z at promoter proximal nucleosomes, as both ‘faces’ of the nucleosome could serve as viable substrates. However, loss of Bre1 did not have a dramatic impact on H2A.Z distribution under normal growth conditions. One possibility is that our basic chromatin immunoprecipitation analyses lack the sensitivity to detect an increase from 1 to 2 copies of H2A.Z per nucleosome, as the immunoprecipitation efficiency may not change substantially. Alternatively, depletion of H2A.Z on the nucleosomal face occupied by H2B-K123ub may be influenced by additional factors, such as RNAPII elongation, masking an impact of H2B-K123ub loss.

Although loss of Bre1 does not lead to a dramatic change in the steady state RNA pool, *bre1* strains show negative genetic interactions with a variety of mutants that disrupt other chromatin regulators (e.g. SWR1C, RSC, Gcn5), suggesting that it does play a key, albeit redundant role in an essential cellular process^38^. H2B-K123ub has been shown to be key for cellular roles other than transcription, in particular, DNA replication ^11,12^. H2B-K123ub is required for recovery from replication stress, and it has been shown to coordinate the intra-S phase checkpoint and nucleosome assembly^11,12,39^. Here we found that Bre1 is also key for a re-distribution of H2A.Z when cells are exposed to high levels of the replication stress agent, hydroxyurea. As cells slowly progress through S phase in the presence of hydroxyurea, H2A.Z re-distributes from promoter proximal nucleosomes to gene bodies. This re-distribution appears linked to fork progression, as the new genic distributions are more highly enriched at genes adjacent to early firing replication origins, compared to genes near late origins.

It is not clear what promotes H2A.Z re-distribution during replication stress or what function this plays. One possibililty is that H2A.Z may aberrantly accumulate over coding regions at stalled forks, and previous work has indicated that fork stalling is less efficient in the absence of Bre1^9^. We note however, that loss of Bre1 does not have a marked impact on the accumulation of cells in S phase during our HU treatment (Figure S3). Alternatively, a more global distribution of H2A.Z may protect stalled forks from collapse. Such a fork protection role might function in concert with the high levels of H3-K56ac that is found on newly assembled nucleosomes following fork passage, a mark that promotes intrinsic nucleosome dynamics, as well as H2A.Z deposition and removal ^40,41^. A potential role in fork protection may provide an explanation for the synthetic lethality of *bre1 swr1* double mutants.

## MATERIALS AND METHODS

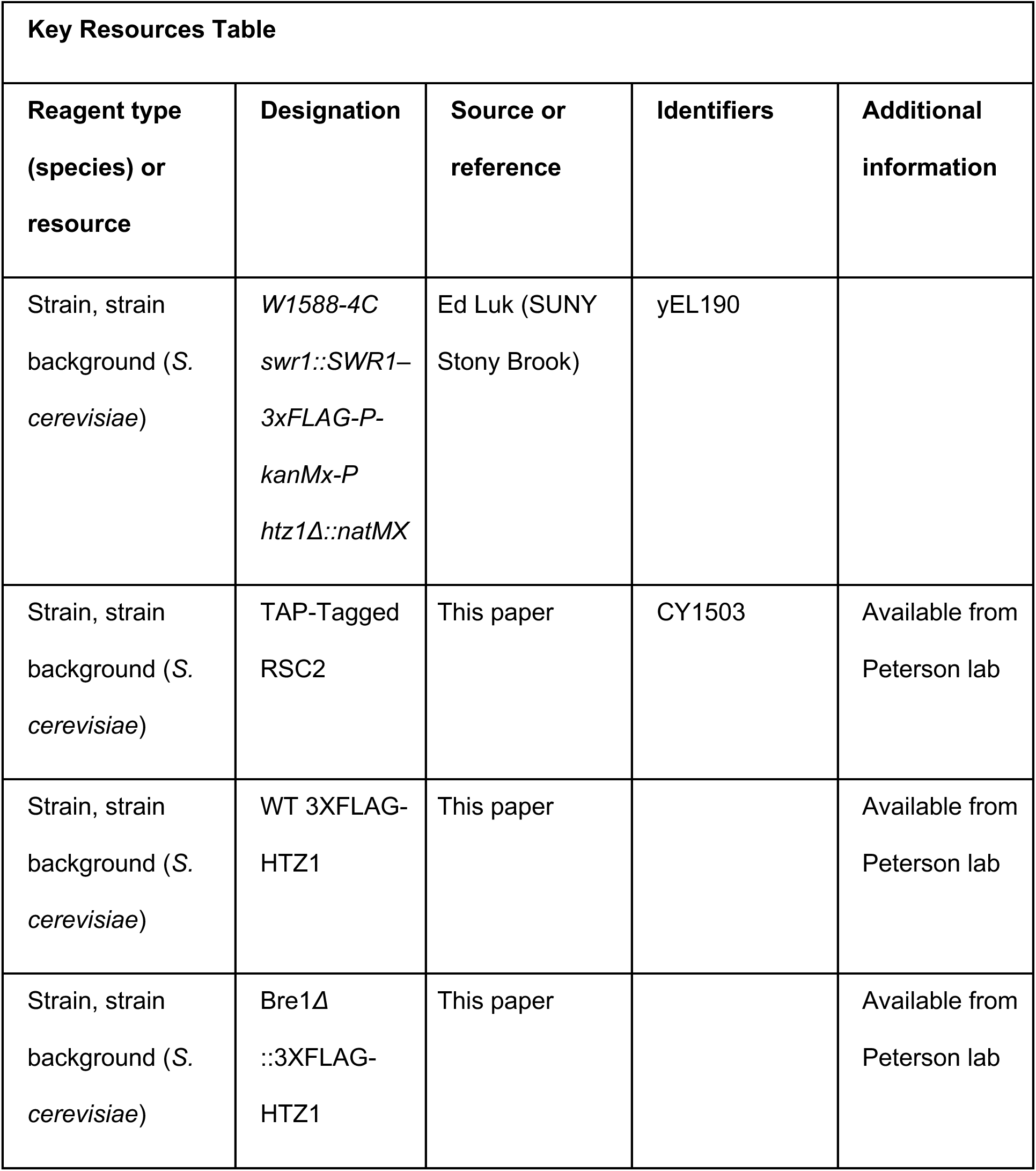
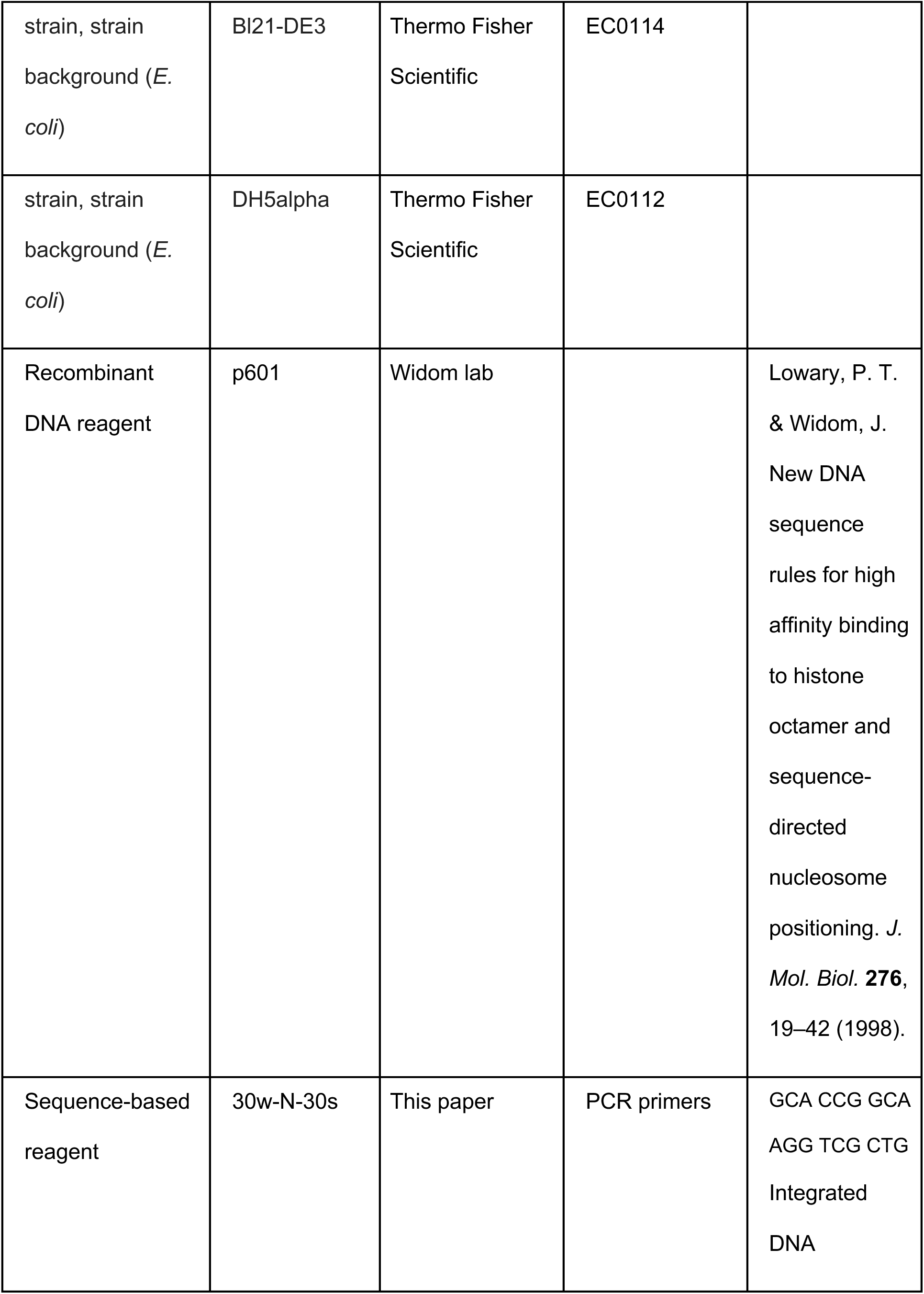
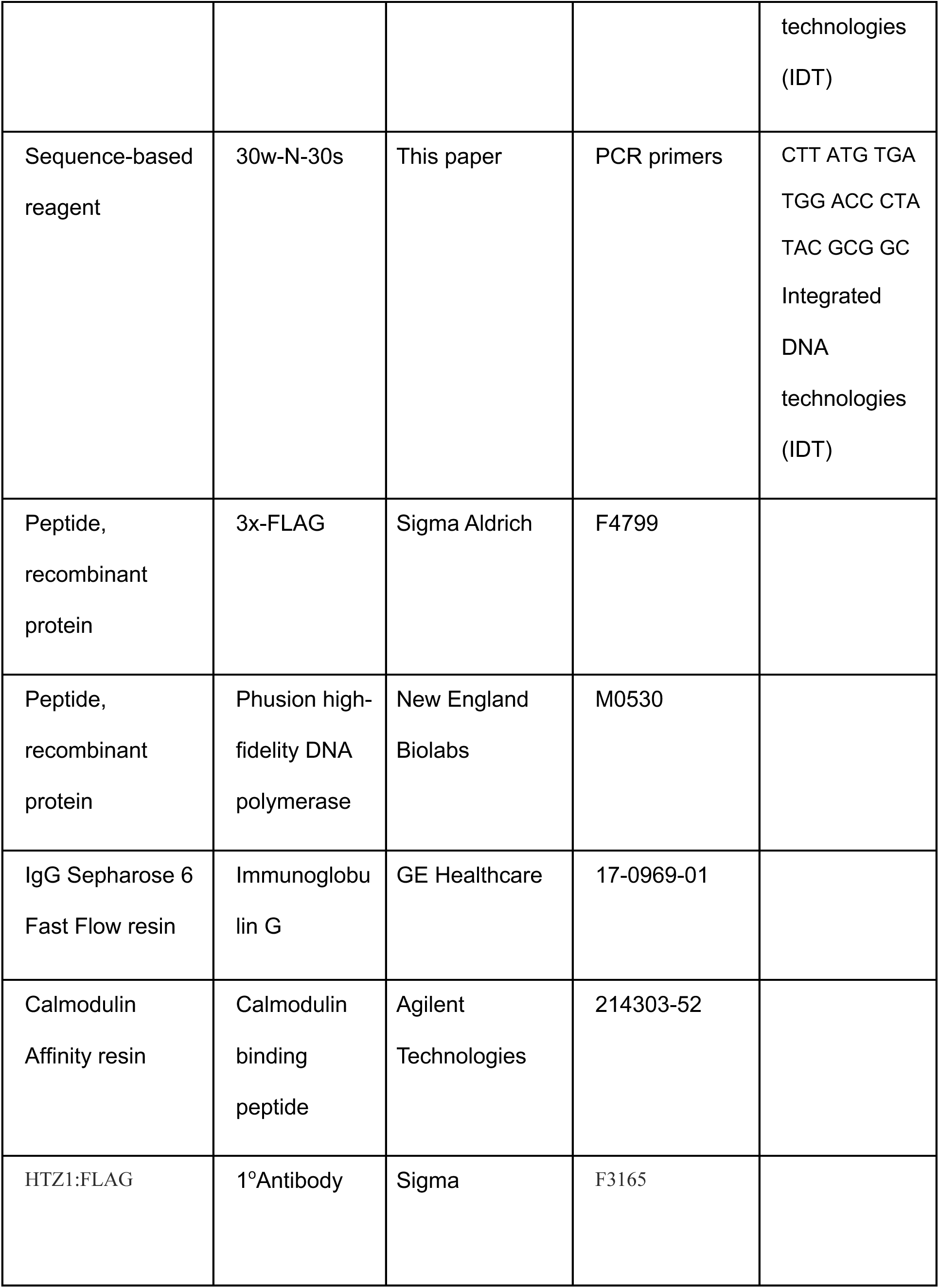
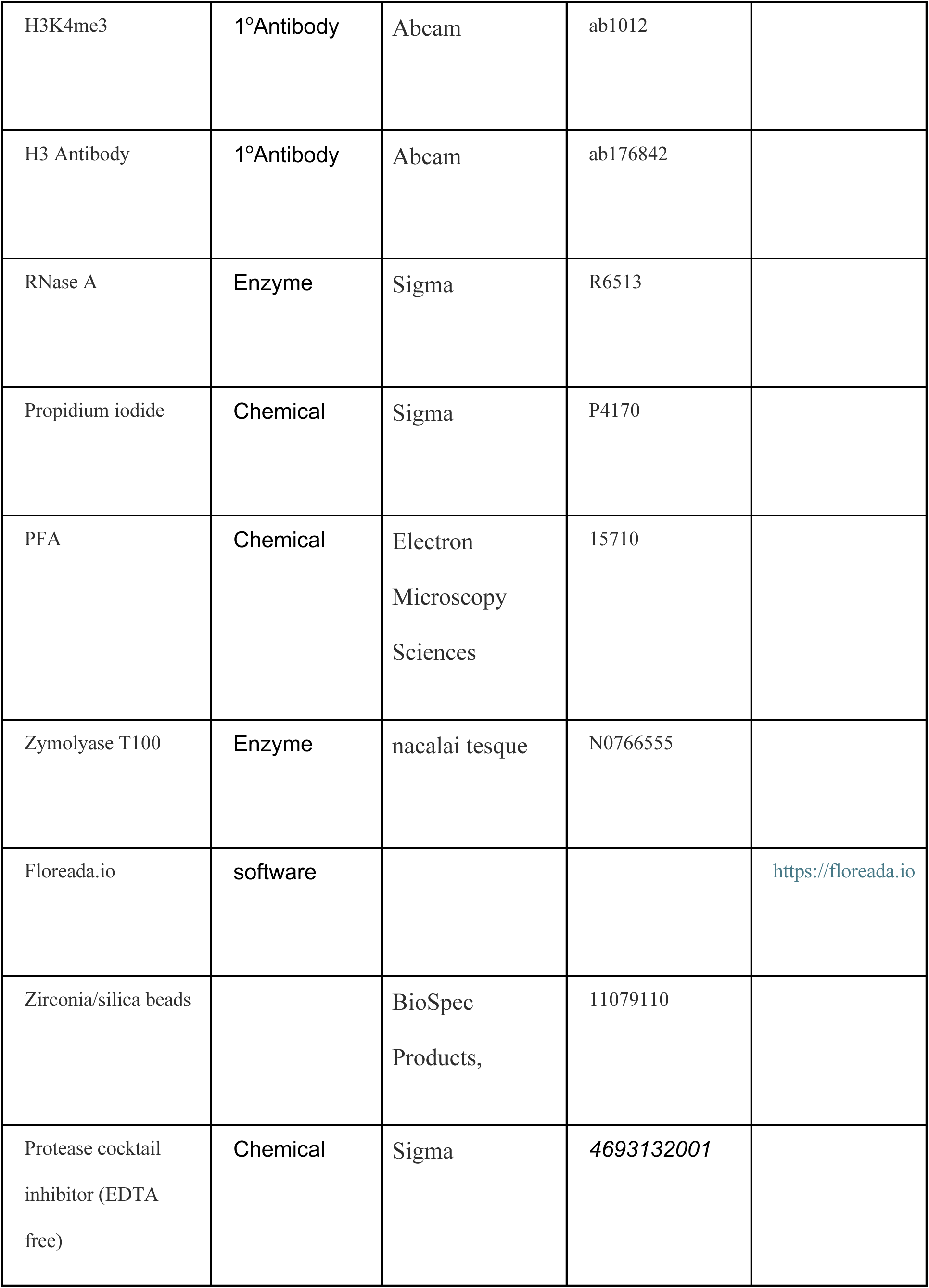
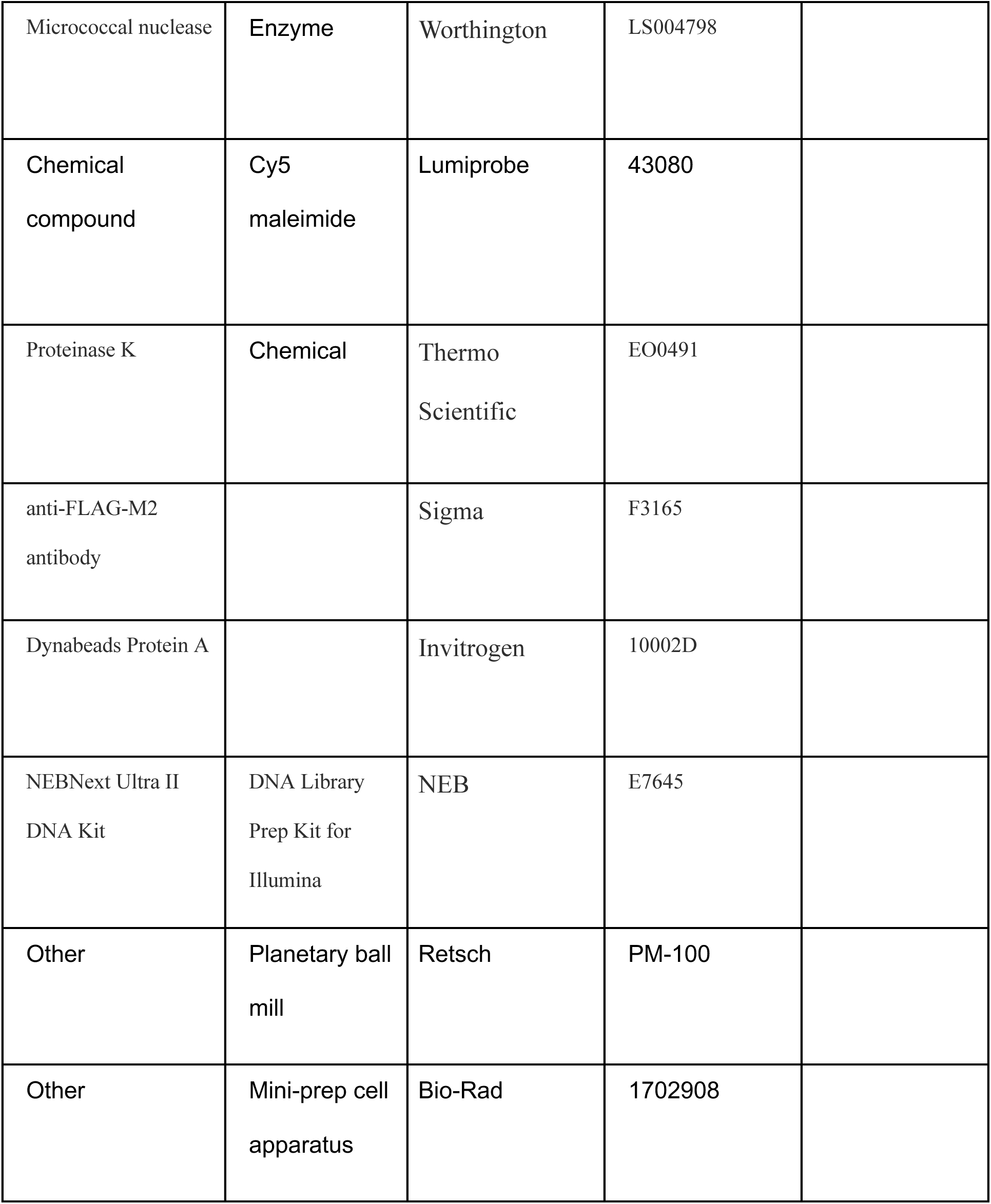

### Yeast strains and culture conditions

The yeast strain *W1588-4C swr1::SWR1– 3xFLAG-P-kanMx-P htz1Δ::natMX* (yEL190) was a gift from Ed Luk (SUNY Stony Brook). The *bre1*Δ** deletion strain was purchased from Euroscarf. For ChIP assays, Htz1 was 3xFlag-tagged in both WT and *bre1*Δ** strains by a PCR-based strategy.

### Protein purification for *in vitro* assays

#### Histones

Histones were prepared as described previously^43^. Histones H2A, H2B, H2B-K123 substitution derivatives, and H2A.Z were from *Saccharomyces cerevisiae,* and histones H3, H4, from *Xenopus laevis* were expressed and purified from Bl21-DE3 expression cells. We used 3X-Flag-tagged and Cy5-labelled yeast ZB dimers for our exchange reactions, and fluorescent label was conjugated onto an engineered cysteine residue of H2A.Z through maleimide linkage as described previously^44,45^.

#### SWR1C

Swr1-3xFlag yeast strains were grown in 6x2L of YEPD, supplemented with adenine at 30°C until reaching an OD_600_ of 3.0. Cells were harvested by centrifuging and noodles were prepared using syringe into liquid nitrogen and stored at -80°C. SWR1C was purified as previously described^45^ with minor modifications. Noodles were crushed into powder using a PM 100 cryomil with 6 × 1 min cycles at 400 rpm, then stored at -80°C. The cell powder (∼200 mL) was resuspended and thawed by adding 200 mL cold 1x HC lysis buffer ([25 mM HEPES–KOH pH 7.6, 10% glycerol, 1 mM EDTA, 300 mM KCl, 0.01% NP40, 10 mM β-glycerophosphate, 1 mM Na-butyrate, 0.5 mM NaF, 1mM DTT plus 1× protease inhibitors [PI 1mM PMSF, 1mM benzamidine, 0.1 mg/mL pepstatin A, 0.1 mg/mL leupeptin, 0.1 mg/mL chymostatin,, added freshly)]. Lysate was cleared by centrifugation at 35000rpm for 3 hours in a Ti45 rotor at 4°C. Whole cell extract was transferred to a 250 mL conical along with anti-Flag M2 agarose resin preequilibrated with 1x HC lysis buffer (1 mL bed volume) and nutated at 4°C for 3 hours. Resin and extract mixture was transferred to an econopac column and after flow through of extract, resin was washed with 4x 25 mL B-0.5 buffer (25 mM <u>HEPES</u>, pH 7.6, 1 mM EDTA, 2 mM MgCl_2_, 10 mM β-glycerophosphate, 1 mM Na-butyrate, 0.5 mM NaF, 500 mM KCl, 10% glycerol, 0.01% NP40 and 1x PI) followed by 3x 10 ml B-0.1 buffer (25 mM <u>HEPES</u>, pH 7.6, 1 mM EDTA, 2 mM MgCl_2_, 10 mM β-glycerophosphate, 1 mM Na-butyrate, 0.5 mM NaF, 100 mM KCl, 10% glycerol, 0.01% NP40 and 1x PI). Elution of SWR1C was performed by incubating the resin on nutator in 1 mL B-0.1 with 0.5 mg/mL recombinant 3xFlag peptide (Genscript) for 1 hour twice in series. The eluants were combined and concentrated using a 50 kDa cutoff Amicon Ultra-0.5 mL centrifugal filter (Millipore). After concentrating down to ∼150 μL, 3x Flag peptide is removed by serial dilution with fresh B-0.1 four more times in the concentrator. The subsequently concentrated SWR1C was aliquoted, flash frozen, and stored at −80°C. SWR1C concentration was determined by SDS-PAGE using a BSA (NEB) standard titration followed by SYPRO Ruby (Thermo Fisher Scientific) staining and quantification using ImageQuant 1D gel analysis.

#### RSC

RSC2-TAP yeast strain was grown in 3x2L of YEPD, until reaching an OD_600_ of 4.0. Cells were harvested by spinning at 3500 rpm in a centrifuge at 4 °C. The cell pellet was passed through a 50ml syringe into liquid nitrogen and noodles were prepared and stored at -80°C. RSC was purified as previously described with minor modifications. Noodles were crushed into powder using a PM 100 cryomil with 6 × 1 min cycles at 400 rpm, then stored at -80°C. The cell powder (∼120 mL) was resuspended and thawed by adding 120 mL cold 1x Lysis Buffer – E Buffer with protease inhibitors ([20mM HEPES, pH7.4, 350mM NaCl, 10% glycerol, 0.1% Tween-20 plus 1× protease inhibitors [PI 1mM PMSF, 1mM benzamidine, 0.1 mg/mL pepstatin A, 0.1 mg/mL leupeptin, 0.1 mg/mL chymostatin,, added freshly]) in a 1L beaker with a stir bar at 4°C. Lysate was cleared by centrifugation at 35000rpm for 3 hours in a Ti45 rotor at 4°C. Whole cell extract was transferred to a 250 mL conical along with 200∼350ul IgG resin equilibrated with 1x E buffer and nutated at 4°C for 2.5 hours. Transfer the resin and extract mixture to an econopac column and after flow through of extract, resin was washed with 3x 5mL of E buffer. After washing resin was incubated with 5ml of E buffer supplemented with 300 units TEV protease and nutated at 4°C for overnight. Next day, the mixture was added to a gravity column and flow through was collected. The flow through from TEV cleavage was supplemented with CaCl_2_ up to 2mM and incubated and nutated for 2 hours at 4°C with preequilibrated Calmodulin resin. The mixture was loaded onto a gravity column and flow through was extracted and the beads were washed with 2X 10mL E buffer. The bound RSC was eluted by adding 3mL E buffer containing 10mM EGTA. Collected eluant was concentrated using a 50 kDa cutoff Amicon Ultra-0.5 mL centrifugal filter (Millipore). After concentrating down to ∼150 μL, the eluted RSC was dialyzed against 500mL E buffer with PMSF, 50uM ZnCl_2_ and 1mM DTT for 3 hours at 4°C. The subsequently concentrated RSC was aliquoted, flash frozen, and stored at −80°C. RSC concentration was determined similarly as SWR1C by SDS-PAGE using a BSA (NEB) standard titration followed by SYPRO Ruby (Thermo Fisher Scientific) staining and quantification using Image Quant 1D gel analysis.

### Nucleosome reconstitution

Unmodified histones were reconstituted into yeast H2A/H2B-Xenopus H3/H4 octamers and yeast H2A.Z/H2B dimers as previously described^46^. The purified octamers and dimers were diluted 1:1 with freeze buffer (10 mM Tris-HCl, pH 7.4, 2 M NaCl, 40% glycerol, 1mM DTT), flash frozen in aliquots, and stored at −80°C for nucleosome reconstitution and downstream assays. Mononucleosomes were reconstituted on DNAs containing the Widom 601 positioning sequence with 31bp linker on both sides (centrally positioned) or 70bp linker on proximal side and 0bp on distal side (end positioned) by salt gradient dialysis as previously described.We have previously demonstrated there is no difference in SWR1C eviction activity on full yeast or hybrid yeast-xenopus nucleosomes^45^.

The ubiquitinylated yeast H2B-K123 analog [yH2B K123ub] was produced by treating yH2B(K123C) with 1,3-dibromoacetone (DBA) and hexahistidine-tagged ubiquitin^47^ and purified by Talon metal affinity and Source S cation-exchange chromatography. This yH2B-K123ub protein and purified yeast H2A were mixed in equimolar ratios in 6 M guanidine-HCl and refolded by dialysis into 10 mM HEPES pH 7.5, 100 mM NaCl, and the yH2A/yH2B K123ub dimer then purified by Source S cation-exchange chromatography. Purified Xenopus histone tetramer and the yH2A/yH2B-K123 ub histone dimer were reconstituted with 207 bp Widom 601 DNA (31 bp flanking each side of the central 145 bp 601 nucleosome positioning sequence) and purified by Source Q anion-exchange chromatography to produce the yH2B-K123ub nucleosomes.

### Nucleosome dimer exchange assays

Gel based dimer exchange assays were performed as done previously with minor modifications^48^. 10nM mononucleosomes (H2B-K123, H2B-K123ub, H2BK123 substitution derivatives), were incubated with 30nM SWR1C and 60nM 3X-Flag tagged H2A.Z/H2B dimers in a 70 μL dimer exchange reaction buffer supplemented with 0.1 mg/mL BSA and 1mM or 10mM DTT. The higher level of DTT had no impact on H2A.Z deposition. The reactions were started by adding 1mM ATP and performed at room temperature. 10 μL of the reaction was taken at each time point and quenched with 1 μg of plasmid DNA to separate SWR1C and dimers from the nucleosome. Each time point was stored on ice until last sample was quenched. Each time point was loaded onto a 6% 0.5x TBE Native PAGE gel (29:1 Acrylamide/Bis) and electrophoresed for 90 min at 120V. Gels were stained with SYBR gold and imaged on a GE Typhoon.

Alternatively, H2B-K123 and H2B-K123ub nucleosomes (10nM) were incubated with (30nM) SWR1C in a 70ul reaction mixture similarly, as above except 3X Flag-tagged H2A.Z/H2B dimers were replaced with Cy5 labelled H2A.Z/H2B dimers. Time points were collected and stored on ice until loaded on 6% PAGE and electrophoresed for 90 min at 120V. Gels were visualized and imaged on a GE Typhoon using Cy5 channel and then stained with SYBR gold and imaged on a GE Typhoon in SYBR gold channel. Band intensity of Cy5 was calculated using Image Quant 1D gel analysis at each time point and plotted against time.

### Restriction Accessibility assays

RAA were performed at 30°C, in a 60 ul reaction with 5nM of RSC, 20nM 31N31 nucleosomes (WT or H2B-K123ub nucleosomes), and 10 units of *Hha*I in a remodeling buffer (20mM HEPES pH 7.8, 3mM MgCl_2_, 0.08% NP-40, 1.7% glycerol, 0.2mM PMSF, 2mM DTT added freshly). Reactions were initiated by adding 1mM ATP. 10ul aliquots were taken at each time point and were stopped with DNA loading dye containing SDS. Each time point was stored on ice until end and loaded on to 5% native PAGE and electrophoresed for 40 min at 100V. Gels were stained with SYBR green and imaged with GE Typhoon. The intensities of cut and uncut bands were measured using Image Quant 1D gel analysis and the ratio was plotted against time.

### Extraction of histone proteins

Yeast pellets were resuspended in 0.4 M H_2_SO4 and disrupted with 1.0 mm zirconia/silica beads (BioSpec Products, 11079110) on vortex for 1 h. Cell debris and acid-insoluble proteins were pelleted by centrifugation at 16,000 g for 5 min at 4 °C. The histone proteins contained in the supernatant were precipitated by dropwise addition of trichloroacetic acid to a final concentration of 33% and incubation on a rotating wheel at 4 °C for 1 h. Precipitated histones were centrifuged at 16,000 g for 10 min at 4 °C, washed twice with acetone and resuspended in ddH_2_O. The presence of HTZ1:FLAG (Sigma, F3165), H3K4me3 (Abcam, ab1012) and H3 (Abcam, ab176842) was assessed by Western blot.

### Cell cycle arrest

S-phase arrest was performed by exposure of log-phase growing WT and *bre1Δ* cells (initial O.D. 0.2) to 200 mM hydroxyurea in YPD medium for 1 h or 2 h 30 min. The cell cycle progression was assessed by assaying DNA content using propidium iodide incorporation. Briefly, cell pellets were fixed in 70% ethanol and subsequently permeabilized in a solution of 0.5% Triton X-100 and 0.1 N HCl in PBS. RNA was degraded by incubating cell pellets in a PBS solution containing 250 µg/ml RNase A (Sigma, R6513) for 2 h at 37 °C. Cells were resuspended in PBS and DNA was stained by addition of propidium iodide (Sigma, P4170) to a final concentration of 4 µg/ml. Flow cytometry data was acquired on a FACSCelesta (BD Biosciences) using a the FACSDiva software with a custom gating panel. FCS files were analyzed with the free online software Floreada.io (https://floreada.io) and the PE-CF594-A parameter, corresponding to PI, was used to fit a cell cycle model using the “Cell Cycle” function. Data from three independent biological replicates was summarized and visualized using a custom R script. Statistical significance was assessed using a t-test.

### Chromatin fractionation

Yeast pellets were resuspended in 0.76 M sorbitol containing 1 mg/ml Zymolyase T100 (nacalai tesque, N0766555) and 25 mM ß-mercaptoethanol for 30 min at 30 °C to allow for cell wall digestion. Spheroplasts were isolated by centrifugation at 3000 g for 5 min, washed once with 0.85 M sorbitol and lysed by resuspension in NP-S buffer (10 mM Tris-HCl pH 7.4, 50 mM NaCl, 5 mM MgCl_2_, 1 mM CaCl_2_, 1 mM ß-mercaptoethanol, 0.5 mM spermidine, 0.075% NP-40) supplemented with protease inhibitors. The insoluble chromatin fraction was isolated by centrifugation at 20,000 g for 10 min at 4 °C. The supernatant containing the soluble fraction was clarified by an additional centrifugation step at 20,000 g for 10 min at 4 °C. The pelleted chromatin fraction was washed once in NP-S buffer and then resuspended in a solution of 50 mM Tris-HCl, 4 mM EDTA and 2% SDS to allow for solubilization of the chromatin proteins. To allow complete resuspension, the chromatin fraction was boiled at 95 °C for 10 min and then sonicated in a bioruptor at medium intensity with intervals of 30 s ON, 30 s OFF for 5 min or until the extract was completely clear. Protein extracts from equivalent number of cells were used for SDS-PAGE. The presence of HTZ1:FLAG (Sigma, F3165) and H3 (Abcam, ab176842) was assessed by Western blot.

### Chip-seq

WT and *bre1Δ* cells were grown on YPD until an O.D. of 0.2 and then incubated with and without 200 mM Hydroxy Urea for an additional 2h 30m. Cell pellets from 250 ml of culture were crosslinked with 1% paraformaldehyde (PFA, Electron Microscopy Sciences, 15710) in PBS for 10 min at room temperature. PFA was quenched by addition of 125 mM Glycine and washed twice with PBS. Cell wall was digested by incubating cell pellets in 0.76 M sorbitol containing 1 mg/ml Zymolyase T100 (nacalai tesque, N0766555) and 25 mM ß-mercaptoethanol for 30 min at 30 °C. Spheroplasts were isolated by centrifugation and washed with 0.76 M sorbitol. Cell lysis was induced by resuspending spheroplasts in NP-S buffer (10 mM Tris-HCl pH 7.4, 50 mM NaCl, 5 mM MgCl_2_, 1 mM CaCl_2_, 1 mM ß-mercaptoethanol, 0.5 mM spermidine, 0.075% NP-40) supplemented with protease inhibitors cocktail (sigma). The chromatin insoluble fraction was pelleted by centrifugation at 20000 g for 10 min at 4°C. Chromatin was fragmented by incubation with 200 U/ml of micrococcal nuclease (Worthington, LS004798) in NP-S supplemented with protease inhibitors cocktail at 37 °C for 5 min The digestion was halted by shifting the reactions to 4 °C and adding EDTA to a final concentration of 20 mM. Further chromatin solubilization was achieved by sonication using a Covaris S220 with peak power 105 W, duty factor 2.0 and 200 cycles/burst for 5 min. Cell debris were separated by centrifugation at 10,000 g for 10 min at 4 °C. Soluble chromatin fraction was quantified as follows: a 5% aliquot was resuspended in three volumes of NP-S containing 200 mM NaCl, 20 mM EDTA, and 0.2 mg/ml Proteinase K (Thermo Scientific, EO0491) and incubated at 65 °C overnight to induce protein digestion and reverse crosslinking. DNA was purified using the Phenol-Chloroform method, precipitated with 75 mM sodium acetate in 75% ethanol, resuspended in water and quantified using nanodrop. For each FLAG ChIP experiment, 30 µg of chromatin equivalents were diluted in five volumes of IP buffer (15 mM Tris-HCl pH 8, 150 mM NaCl, 1 mM EDTA 1% Triton X-100) with the addition of 4 µg of monoclonal anti-FLAG-M2 antibody (Sigma, F3165) and 20 µL of Dynabeads Protein A (Invitrogen, 10002D) and incubated overnight at 4 °C on a rotating wheel. Beads-bound chromatin was separated using a magnetic rack and washed twice with IP buffer and twice with high-salt buffer (IP buffer with a final concentration of 450 mM NaCl). Immunoprecipitated chromatin was eluted by resuspending the beads in 1% SDS and 0.1 M NaHCO3 in a thermomixer at 65 °C for 1 h at 1000 rpm. The supernatant containing immunoprecipitated chromatin was de-crosslinked overnight at 65 °C with the addition of 200 mM NaCl. To perform protein digestion, each sample was added 10 mM EDTA, 40 mM Tris-HCl pH 6.5, and 0.2 mg/ml Proteinase K and incubated at 45 °C for 1 h. DNA was purified as described above. Sequencing libraries were prepared using the NEBNext Ultra II DNA Library Prep Kit for Illumina (NEB, E7645) according to manufacturer’s instructions. Libraries were paired-end sequenced on an Illumina NextSeq 500 platform.

### ChIP-seq data analysis

Paired-end ChIP-seq reads were adapter-trimmed using trim-galore (v0.6.10) and aligned to the sacCer3 genome using Bowtie 2 (v2.5)^49^ with the parameters -t -q -N 1 -X 1000 --no-mixed --no-discordant --no-unal --local --very-sensitive-local. PCR duplicates were removed using the rmdup command in samtools (v1.16.1). The deepTools suite (v3.5.6)^50^ was used to generate genome coverage files, compute the average of biological replicates and compute gene metanalyses to produce heatmaps and metagene profiles. Peak calling was performed using MACS2 (v2.2.7.1) and reproducible peaks between biological replicates were identified using IDR (v2.0.2)^51^. Peaks were annotated onto genomic features using the ChIPseeker package (v1.40) in R (v4.2.2), with promoters being defined as the regions going from 250 bp upstream to 50 bp downstream of transcription start sites. A list of 85 early replicating and 85 late replicating genes were manually curated based on their location within 1-2 kb of efficiently firing early or late origins^52^ (Table S1).

## Figure Legends

**Figure S1.**
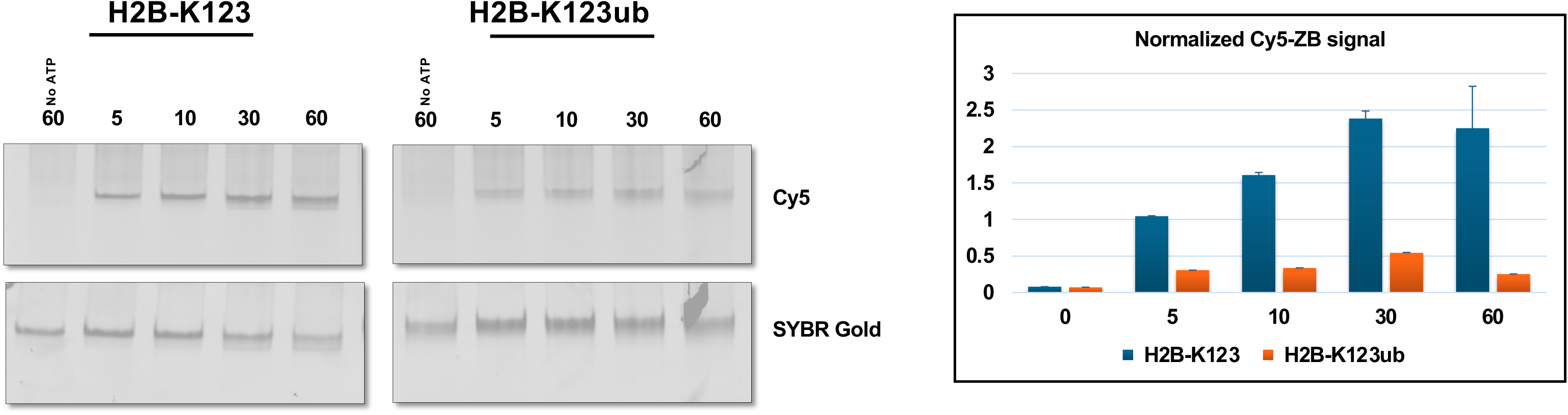
H2B-K123ub inhibits SWR1C-mediated dimer exchange activity. Left panels. Upper panels show dimer exchange activity using Cy5-labelled dimer and H2B-K123 and H2B-K123ub nucleosomes. Lower panels show same gels stained with SYBR gold and imaged with Typhoon imager. Right panel. Quantification of right panels. We note that this assay is prone to nonspecific, ATP-independent association of Cy5-labelled dimers with nucleosomes, leading to background signals.

**Figure S2.**
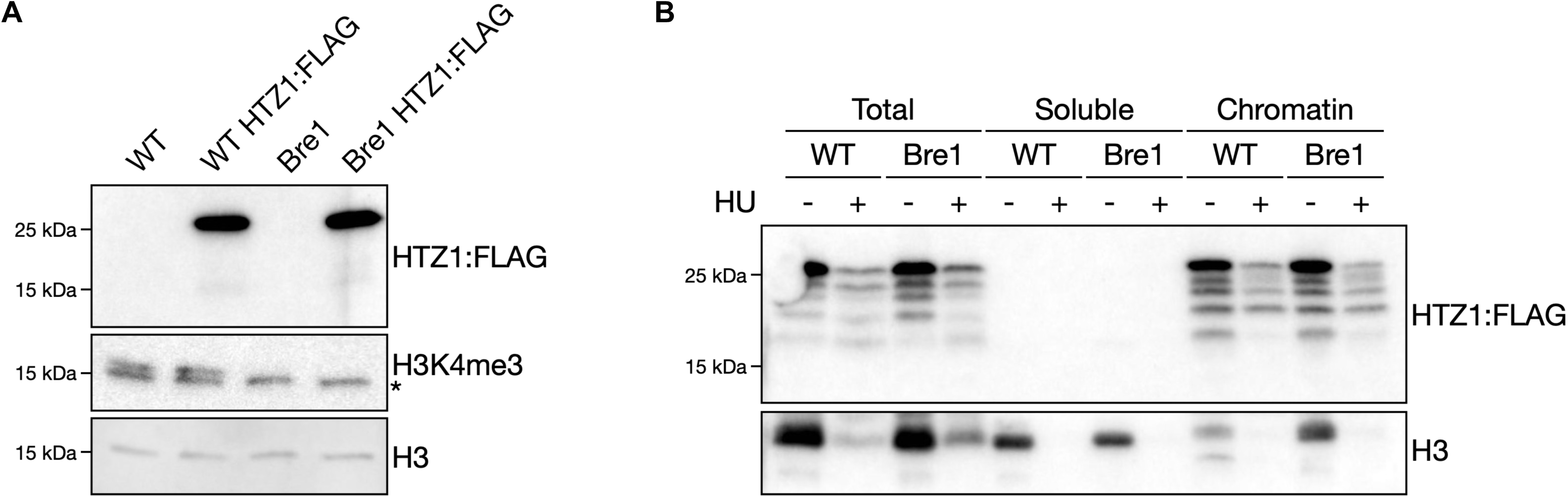
Bulk levels of Htz1 are not altered in *bre1*Δ** cells. a) Western blot of histone extracts from WT and *bre1*Δ** cells expressing or not the Htz1:FLAG construct. The asterisk in the H3-K4me3 blot indicates the band corresponding to H3; b) Western blot of total, soluble, and chromatin fractions of WT *HTZ1:FLAG* and *bre1*Δ* HTZ1:FLAG* cells upon incubation with or without HU.

**Figure S3.**
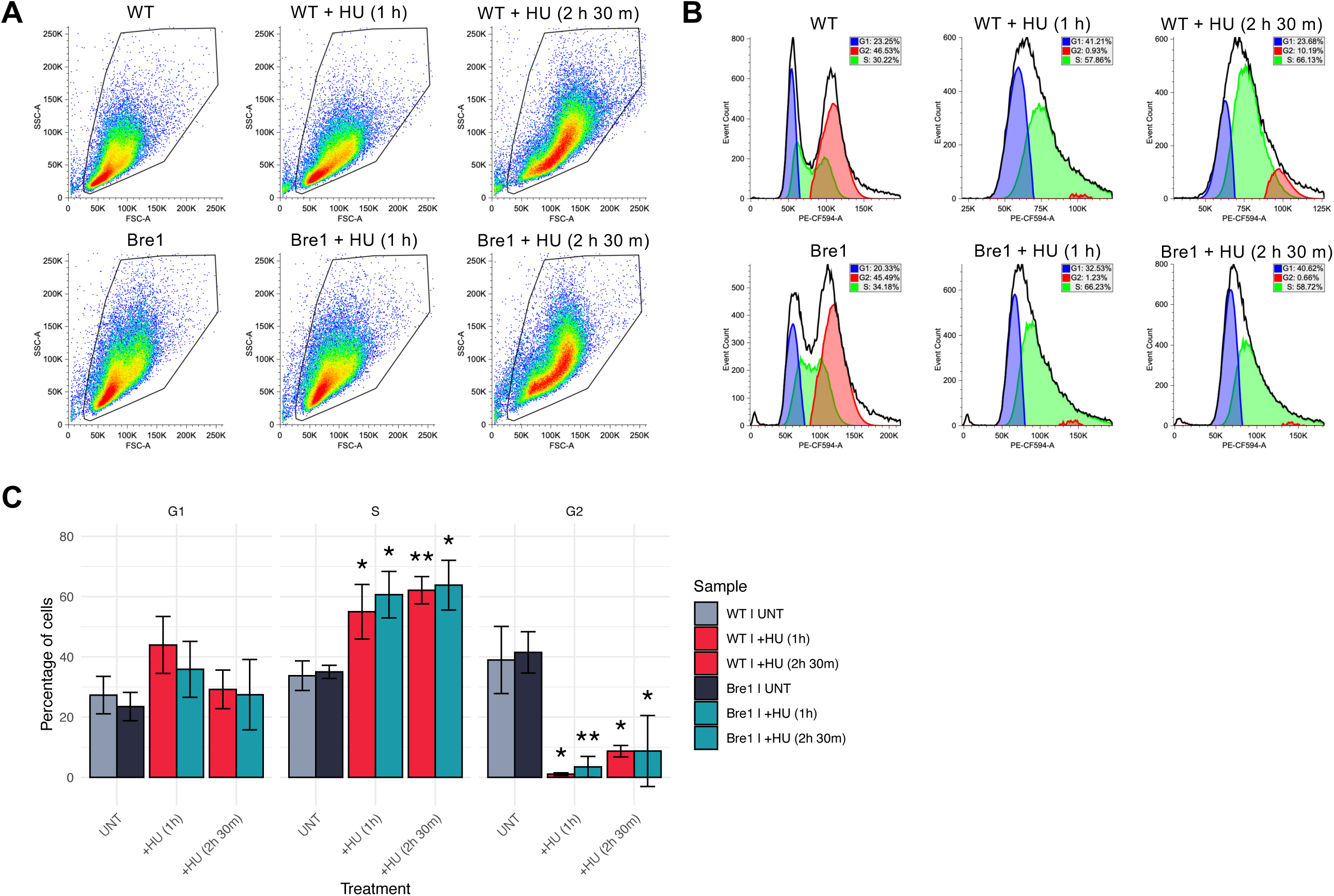
HU induces cell cycle arrest in S-phase. a) Representative example of a FACS plot of the forward and side scatter of WT and *bre1*Δ** cells upon HU treatment; b) Representative example of PE-CF594-A signal intensity, corresponding to incorporated propidium iodide, the blue, red and green shades below the curve indicate the modelled cell cycle phases (G1, G2, and S phase, respectively); c) Quantification of the percentage of cells in each phase of the cell cycle upon HU treatment, as shown in (b). Each bar and error bar represent the mean and standard deviation of three independent biological replicates; Statistical significance of the effect of HU treatment to each phase of the cell cycle was assessed using the t-test (* p < 0.05, ** p < 0.01).

